# V genes in rodents from whole genome sequencing data

**DOI:** 10.1101/011387

**Authors:** David N. Olivieri, Santiago Gambón-Cerdá, Francisco Gambón-Deza

## Abstract

We studied the V exons of 14 rodent species obtained from whole genome sequencing (WGS) datasets. Compared to other mammals, we found an increase in the number of immunoglobulin (IG) V genes in the heavy (IGH) and kappa chain (IGK) loci. We provide evidence for a reduction genes in lambda chain (IGL) locus, disappearing entirely in one of the species(*Dipodomys ordii*). We show relationships amongst the V genes of the T-cell receptors (TR) found in primates, possessing ortholog sequences between them. As compared with other mammals, there is an increase in the number of TRAV genes within rodents. Such an increase within this locus is caused by duplication events involving a few putative V genes. This duplication phenomenon does not occur in the TRBV locus. In those species that underwent an expansion of TRAV genes, we found that they also have a correspondingly larger number of MHC Class I genes. The results suggest that selective pressures have conditioned the expansion of V genomic repertoire the TRA, IGK and IGH loci during the diversification process of rodents.

## 1. Introduction

Antigen recognition in the immune system of vertebrates is carried out by the immunoglobulin (IG) and T-cell receptor (TR) molecules. In these molecules, variable regions exist that are complementary to antigen (Janeway et al., 2005). Immunoglobulin (IG) recognizes antigen directly in soluble form and the antibody-antigen binding site is composed of two NH2-terminal protein chains, called the heavy (IGH) and light chain (IGK and IGL) (Guddat et al., 2000). The interaction region, where recognition takes place, is encoded by V genes. In IGV genes, there are three separate loci in mammals (Wu & Kabat, 1970), one for the heavy chain (IGHV) and two for the light chains (i.e., one for kappa genes (IGKV) and one for lambda (IGLV) genes) (Brack et al., 1978). The constellation of such genes, distributed across these three loci,constitute the germinal immunoglobulins V gene repertoire of a specie. Moreover, during the development of an individual, recombination with D and J genes and processes of somatic mutations work together to condition the recognition capabilities of the germinal gene repertoire (Tonegawa, 1983; Davis & Bjorkman, 1988).

The most detailed observation of the interaction of an antibody with its antigen has identified three putative regions within the V region that are involved in the contact with antigen(Kabat & Wu,1991; Mian et al., 1991). These interaction sites are referred to as the complementarity determining regions (CDR). Thus, the recognition process involves the interaction of six CDR (three for each chain) with antigen. For each IG chain, two of the CDR are encoded within the V exons, while the third is formed in processes of somatic recombination (Lefranc & Lefranc, 2001a; Lefranc, 2001) (see also: The Immunoglobulin FactsBook Lefranc & Lefranc (2001a); The T cell receptor FactsBook Lefranc & Lefranc (2001b)).

T lymphocytes also recognize antigen, but in denatured form and together with the major histocompatibility complex (MHC) molecules (Davis & Bjorkman, 1988). While the recognition mechanisms are different IG and TR have a similar molecular structure. In the case of TR, antigen-MHC recognition is performed by two chains involving two V regions. As in the case of IG chains, each TR V region has three CDR so that a total of six CDR enter into close contact with the antigen-MHC complex (see the *International ImMunoGeneTics Information System*, http://www.imgt.org Lefranc et al. (2009); Lefranc (2011b), IMGT/GENE-DB Giudicelli et al. (2005). Each chain of the TR V genes originate from germline sequences on different loci. Similar D and J gene recombination processes and somatic mutations occur amongst the TR V gene loci, equivalent to those described for IG loci (Janeway et al., 2005).

Vgenextractor (Olivieri et al., 2013) is a bioinformatics tool that obtains 90% of V exon sequences of all loci from whole genome sequencing (WGS) datasets by identifying conserved motifs in the germline sequences. With this program we have obtained tens of thousands of V exon sequences of jawed vertebrates that have been deposited in a public and freely accessible repository, *vgenextractor.org*. Recently, we confirmed the results of Vgenextractor with an alternative approach based upon the random forest method that makes no prior assumptions about motifs. By training the random forest with known V exon annotations, predictions provides a deeper probabilistic exploration of the WGS dataset than was possible with our previous approach.

Using the data obtained from our software, we recently studied the V gene repertoire in primates (Olivieri & Gambon-Deza, 2014). We show evolutionary patterns for IG V genes (common processes of birth and death) and positive selection pressure that gives rise to V gene conservation in the TR loci. In particular, we identified 35 TRAV and 25 TRBV conserved genes across primate species. Due to the evolutionary implications of these results, we study here whether similar phenomena occur in the order Rodentia, by analyzing available WGS from representative species.

## 2. Material and methods

### The Genome Data

We studied sequences extracted from 14 Rodent WGS genome assemblies (acquired from the NCBI) and deposited in our repository, *vgenextractor.org*. Salient information about the genome assemblies studied is provided in Table 1. While the WGS datasets are in various stages of maturity, the average contig N50 for the Rodent species studies was 38000 with an average coverage of 100× for Illumina sequencing, and 6× for the Sanger sequencing method.

**Table 1:**
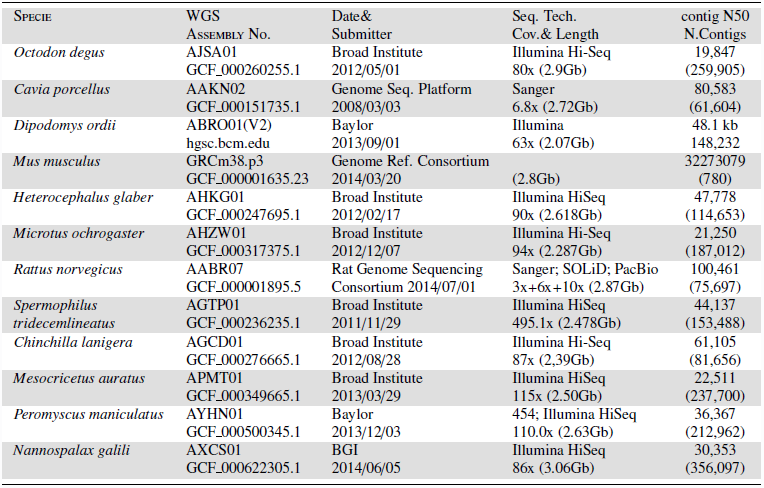
WGS Data for the 14 rodent species

The WGS used are provided in Figure 1 of the phylogenetic tree. A summary of the accession abbreviated numbers for each species is the following: *Octodon degus* (AJSA01), *Cavia porcellus* (AAKN02), *Dipodomys ordii* (ABRO01), *Mus musculus* (AAHY01), *Heterocephalus glaber* (AHKG01), *Ochotona princeps* (ALIT01), *Microtus ochrogaster* (AHZW01), *Oryctolagus cuniculus* (AAGW02), *Rattus norvegicus* (AABR06), *Spermophilus tridecemlineatus* (AGTP01), *Chinchilla lanigera* (AGCD01), *Cricetulus griseus* (AFTD01), *Jaculus jaculus* (AKZC01), *Mesocricetus auratus* (APMT01), *Peromyscus maniculatus* (AYHN01) and *Nannospalax galili* (AXCS01).

**Figure 1:**
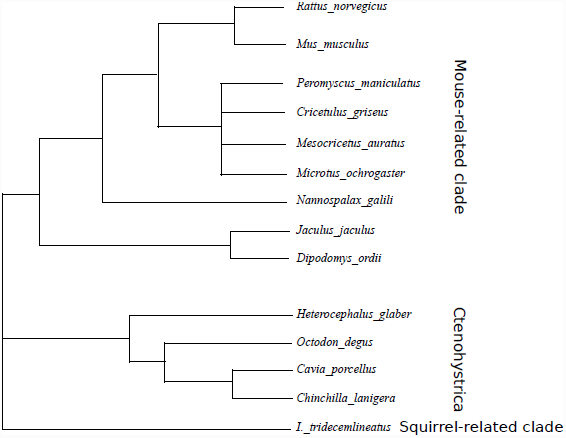
Phylogenetic trees of species in the Rodentia order considered in this study. The corresponding abbreviated WGS code and assembly are indicated for each species.

### V exon identification with Random Forest

We recently developed a random forest technique for V gene identification (Olivieri & Gambon-Deza, 2014), which serves as an alternative and separate validation of the Vgenextractor tool. With this method, we predicted homologous V exon sequences by training our model with non-rodent mammalian V exon sequences annotated from the IMGT and those at vgenextractor.org. In our random forest method, the in-frame nucleotide exon sequence was translated to amino acids which was used to derive a 500 element vector with a *physicochemical distance transform* (PDT) (Liu B, 2012). Our multiclass training set consisted of 16829 positively identified V exon sequences (the positive sequences in each locus were: IGHV: 3512, IGKV: 2396, IGLV: 3065, TRAV 4384, TRBV: 2150, TRDV: 600, TRGV: 722) and 51790 random sequences (a background/signal ratio of 3:1, that represent negative background. Thus, the prediction automatically determines the locus based upon a maximal probability measure. Both the Vgenextractor and our random forest prediction tool are available through vgenextractor.org.

### Comparative Analysis

For comparative sequence analysis, we performed alignments with ClustalO (Sievers & Higgins, 2014) in the SEAVIEW (Gouy et al., 2010; Sievers & Higgins, 2014) environment and constructed trees with FastTree (Price et al., 2010), Figtree (Rambaut) and Jalview software (Waterhouse et al., 2009). We also developed our own python scripts that used the Biopython and Dendropy (Sukumaran & Holder, 2010) libraries for solving specific phylogenetic problems. We also used the matplotlib library (Hunter, 2007), particularly to obtain ‘radar’ plots to illustrate maximal clade membership relationships.

### MHC Correlation studies

We developed a software tool to explore whether a V gene expansion in a species corresponds to an expansion in the number of MHC genes. In particular, we developed a random forest based algorithm to identify MHC class I (MHC-I) and class II (MHC-II) gene sequences from the WGS datasets of Rodents. From the two sets, we determined whether a positive correlation exists based upon standard statistical measures.

For the case of MHC-I, we obtain the three main exons (EX2, EX3 and EX4, following IMGT G-Domain and MHC nomenclature (Lefranc et al., 2005; Robinson et al., 2011)). In the case of MHC-II, we obtain the two main exons (EX2 and EX3) for both the alpha and beta chain genes. For each constituent MHC exon, we trained separate random forest models based upon non-rodent annotated sequences available at the IMGT/HLA database (Robinson et al., 2011). However, because there is limited annotated sequences for a range of species, we also developed a simple *bootstrapping program* that identifies exons using specific conserved amino acid motifs identified by comparing annotated species. In this way, we obtained sufficient training data for the random forest. Next, we transformed each in-frame amino acid translated exon sequence to a 500 element vector using the *physicochemical distance transform* (PDT) (Liu B, 2012) that captures the positionally dependent physicochemical properties of the sequence.

To obtain MHC-I and MHC-II from the WGS data, we first pre-selected contigs likely to contain MHC exons using a TBLAST (Tatusova & Madden, 1999) query with human MHCs and with a large threshold (e¡10). From the set of WGS contigs, our algorithm identifies potential exon sequences by first delineating all sequence intervals between an exon start *AG* and an exon stop *GT* filtered by a minimum and maximum size (e.g., having the necessary nucleotide size for producing an exon of 92 amino acids). Next, the set of valid sequence intervals are translated in-frame to amino acids. If no stop codon is found within the exon, it is transformed, as in the training set, to a 500 element (PDT) vector (Liu B, 2012). Each putative exon sequence is tested against each of the trained random forest exon models (ie., EX2, EX3, and EX4 for MHC-I, and EX2 and EX3 for case MHC-II). In this way, the candidate exons are classified into either one of the exons by a maximum probability score.

Functional MHC molecules must maintain a certain intron/exon ordering. Thus, in the final steps of the algorithm, valid MHC molecule are those that are constrained to the correct exon-intron tandem structure, dependent upon the MHC type (ie., MHC-I or MHC-II). From the list of candidate exons that have a random forest homology score ¿50% in any one of the categories, we also require that the order is precisely maintained. The implementation of our algorithm as well as the MHC sequences obtained from the WGS datasets are available at *vgenextractor.org*.

## 3. Results

Rodents correspond to mammals that are characterized by the presence of continuous growth of incisors. Of all mammal species, 40% are rodents. We studied V exon sequences extracted from the WGS of 14 different rodent species from the datasets deposited at the NCBI repository. The phylogenetic tree of the Rodent order for the species studied in this work is shown in Figure 1 following molecular classification studies (Farwick et al., 2006; Churakov et al., 2010). This grouping distinguishes three main evolutionary clades of rodents: Squirrel-related, Ctenohystrica and Mouse-related.

The translated V exon sequences of these species were obtained using Vgenextractor and independently confirmed with our random forest method (See Methods section). We have shown that our program obtains ¿90% of all V exon sequences in both the IG and TR loci (Olivieri et al., 2013). The number of genes found per locus is given in Table 2. For all species, there are more IGK than IGL genes, and in particular, there are few IGLV exons in the mouse-related clade. In *Dipodomys ordii* no V genes were detected in the IGL locus (confirmed in two different draft versions of the WGS datasets, ABRO01 here was a preliminary kangaroo rat assembly using 7.4 million (2.5×) sanger sequenced reads from Broad Institute and 2.8Gb The final assembly has a size of 2.07Gb with contig N50 of 48.1 kb and scaffold N50 of 11.3Mb *ftp://ftp.hgsc.bcm.edu/Dipodomys_ordii/genome_assemblies/kangaroo_rat.20130901.contigs.fa*, consisting of approximately 3.4 billion Illumina reads with an accumulated coverage ¿100×, while in *J. Jaculus* only one IGLV gene was detected. To the contrary, rodents have a large number of V exons for the heavy chain (IGH) and the kappa chain (IGK) loci. This increase, as compared to other species, appears to be accompanied by a corresponding increase in the number of TRA V exons. In the case of *R. norvegicus*, there are 146 IGHV genes, 174 IGKV and 198 TRAV genes. This expansion in V exon number in *R. norvegicus* is not a general trend manifest in all loci, since the number of IGLV and TRBV genes is relatively low. The elevated number of TRD V genes may be due to the same expansion process witnessed by the TRA V genes, since these exons share the same chromosome region.

**Table 2:** Distribution of V-genes amongst the IG and TR loci.

| SPECIES | IGHV | IGLV | IGKV | TRAV | TRBV | TRDV | TRGV | ALL |
| --- | --- | --- | --- | --- | --- | --- | --- | --- |
| **SQUIRREL-RELATED** |  |  |  |  |  |  |  |  |
| *I. tridecemlineatus* | 12 | 54 | 131 | 76 | 30 | 7 | 12 | 322 |
| **CTENOHYSTRICA** |  |  |  |  |  |  |  |  |
| *H. glaber* | 22 | 13 | 35 | 40 | 15 | 3 | 4 | 132 |
| *C. porcellus* | 97 | 48 | 117 | 71 | 37 | 11 | 11 | 392 |
| *C. lanigera* | 33 | 34 | 65 | 45 | 21 | 8 | 8 | 214 |
| *O. degus* | 125 | 44 | 80 | 62 | 26 | 12 | 16 | 365 |
| **MOUSE-RELATED** |  |  |  |  |  |  |  |  |
| *D. ordii* | 55 | 0 | 86 | 31 | 11 | 1 | 4 | 188 |
| *J. jaculus* | 10 | 1 | 42 | 11 | 21 | 0 | 0 | 85 |
| *N. galili* | 72 | 9 | 97 | 47 | 18 | 7 | 5 | 255 |
| *M. ochrogaster* | 69 | 21 | 46 | 57 | 31 | 3 | 6 | 233 |
| *M. auratus* | 50 | 16 | 64 | 38 | 19 | 1 | 4 | 192 |
| *C. griseus* | 126 | 23 | 55 | 54 | 26 | 3 | 0 | 287 |
| *P. maniculatus* | 105 | 12 | 199 | 174 | 34 | 25 | 5 | 554 |
| *M. musculus* | 109 | 4 | 111 | 93 | 18 | 10 | 7 | 352 |
| *R. norvegicus* | 146 | 15 | 174 | 198 | 20 | 30 | 5 | 588 |

### 3.1. The IGHV genes

In the IGHV locus, previous studies have identified three phylogenetic clans (Kirkham et al., 1992; Lefranc, 2011a). From a phylogenetic analysis of IGHV exon sequences from primates, we have recently identified distinct subclades within each of these pre-established clans. Thus, by using consensus sequences from these primate IGHV subclades, we studied whether there exists evolutionarily related sub-clades amongst the corresponding IGHV exon sequences of rodents.

In Clan-I of primates, we detected three subclades, which we denote I-A, I-B and I-C. No homology was found between the primate consensus sequence of I-A and the rodent sequences of Clan-I, suggesting that one of two possibilities: that an ancestral exon sequences was lost in the speciation from rodents, or that this subclade was generated within Primates. With respect to the subclades I-B and I-C, however, we did find homology.

**Figure 2:**
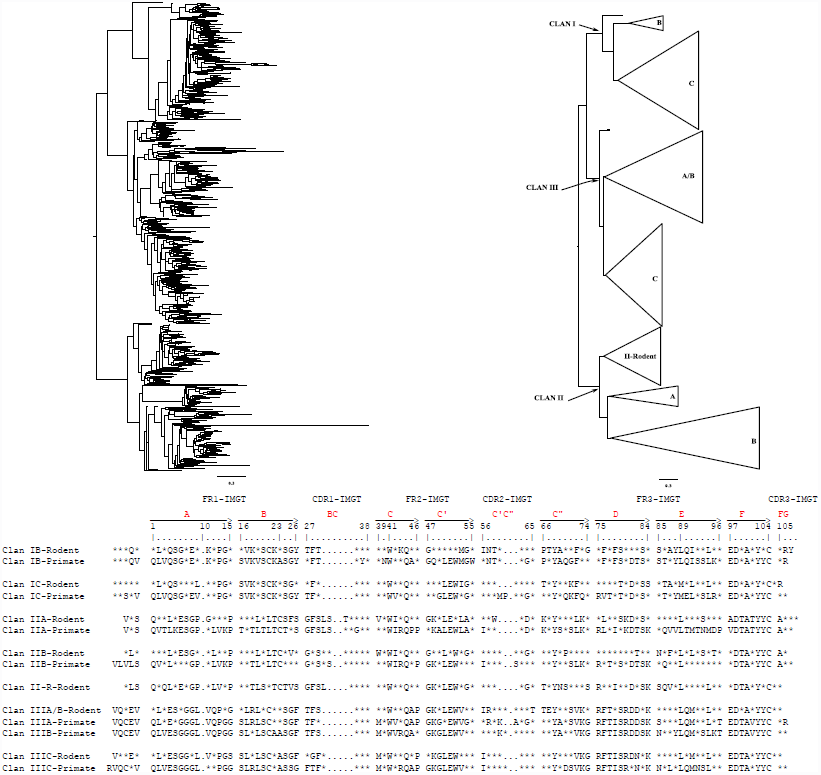
The phylogenetic trees of the AA translated sequences from IGHV exons from 14 rodent species. V exon sequences are obtained from whole genome shotgun (WGS) datasets using the Vgenextractor algorithm (Olivieri et al., 2013) and independently confirmed using a random forest approach. Alignment of the amino acid sequences was performed with clustalΩ (Sievers & Higgins, 2014), tree construction with FastTree (Price et al., 2010) using the WAG matrix, and visualization with Figtree (Rambaut). Left: the tree of all IGHV exon sequences; Right: Clades identified and collapsed. In the bottom part of the figure, the consensus sequences of each clade are shown. The sequences of rodents were aligned with the consensus sequences of primates. The amino acids that are found in more than 90% of the sequences are marked by their letter, while the variable regions are represented by an asterisk (”^*^”)

Similarly, in Clan II of rodents we identified two subclades related evolutionarily to primates, denoted II-A and II-B. In addition, another clade, denoted R (for rodent), that is unrelated to any sequence found in primates. In Clan III of rodents, subclades III-A and III-B of primates are homologous to rodent sequences that form a single clade. Also, in this clade, there is a homologous clade to the III-C subclade found in primates. These observations are summarized in Figure 2 and Table 3.

**Table 3:** Distribution of IGHV exons for each of the defined clans.

|  | CLAN I |  |  | CLAN II |  |  | CLAN III |  |
| --- | --- | --- | --- | --- | --- | --- | --- | --- |
| SPECIE | A | B | C | A | B | R | A/B | C |
| **SKIRREL-RELATED** |  |  |  |  |  |  |  |  |
| *I. tridecemlineatus* | 0 | 0 | 0 | 0 | 6 | 0 | 3 | 3 |
| **CTENOHYSTRICA** |  |  |  |  |  |  |  |  |
| *H. glaber* | 0 | 1 | 2 | 0 | 8 | 2 | 3 | 6 |
| *C. porcellus* | 0 | 0 | 0 | 0 | 25 | 15 | 7 | 53 |
| *C. lanigera* | 0 | 1 | 3 | 1 | 12 | 4 | 6 | 6 |
| *O. degus* | 0 | 6 | 24 | 9 | 18 | 11 | 8 | 49 |
| **MOUSE-RELATED** |  |  |  |  |  |  |  |  |
| *D. ordii* | 0 | 1 | 3 | 4 | 0 | 0 | 3 | 44 |
| *J. jaculus* | 0 | 0 | 0 | 0 | 4 | 1 | 0 | 5 |
| *N. galili* | 0 | 1 | 12 | 1 | 4 | 8 | 3 | 37 |
| *M. ochrogaster* | 0 | 2 | 21 | 1 | 5 | 17 | 7 | 17 |
| *M. auratus* | 0 | 1 | 22 | 0 | 2 | 7 | 9 | 9 |
| *C. griseus* | 0 | 5 | 30 | 8 | 27 | 12 | 19 | 25 |
| *P. maniculatus* | 0 | 4 | 28 | 5 | 25 | 7 | 21 | 14 |
| *M. musculus* | 0 | 4 | 55 | 6 | 8 | 9 | 13 | 14 |
| *R. norvegicus* | 0 | 2 | 25 | 9 | 6 | 41 | 22 | 41 |

From the studies between the consensus 90% sequences of rodents and primates, we have shown the presence of highly conserved amino acids (AA) that have existed in mammalian orders separated evolutionary by more than 70My (Churakov et al., 2010). Such AA conservation is particularly surprising in the CDR sequences, especially in the first three amino acids of CDR1, as shown in Figure 2.

### 3.2. Light chains V genes

As found in primates(Olivieri & Gambon-Deza, 2014), rodents have more V genes in the IGK locus than in the IGL locus. Indeed, of all the mammals we have analyzed for V genes (*vgenextractor.org*), rodents have the highest IGK/IGL ratio. As mentioned previously, in *D. Ordii* no IGL V exon sequences were detected. We have not found the constant region of the lambda chains or any sequence in RNA-seq studies. It is the only mammal described so far that lacks the IGL chains.

**Figure 3:**
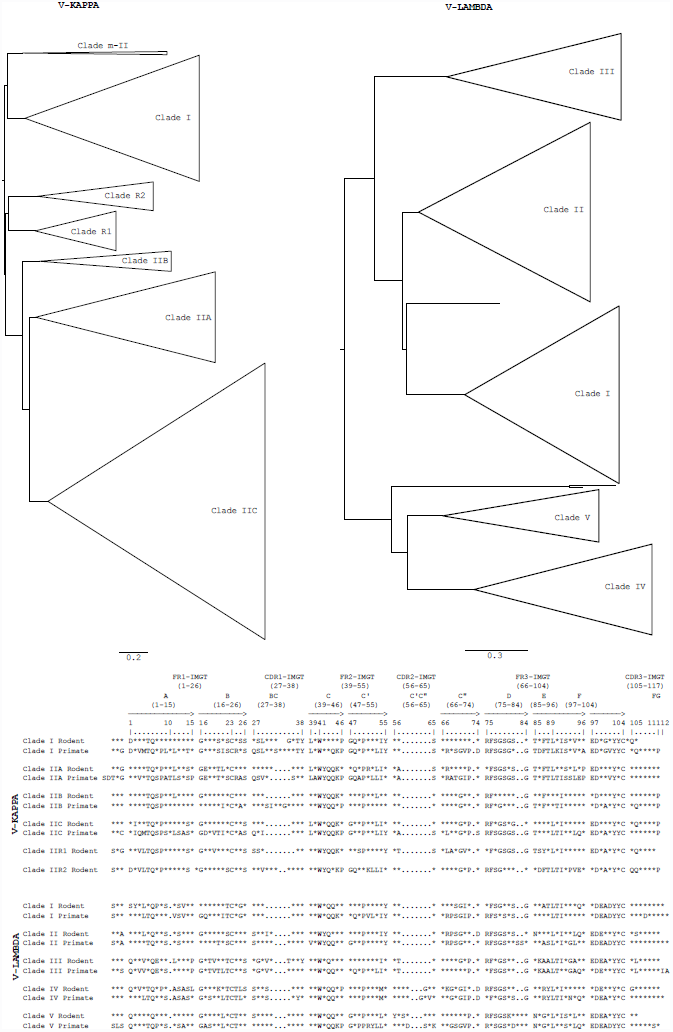
The phylogenetic trees of the AA translated sequences from (IGKV -left-and IGLV -right-) exons from 14 rodent species. V exon sequences are obtained from whole genome shotgun (WGS) datasets using the Vgenextractor algorithm (Olivieri et al., 2013) and independently confirmed using a random forest approach. Alignment of the amino acid sequences was performed with clustalO (Sievers & Higgins, 2014), tree construction with FastTree (Price et al., 2010) using the WAG matrix and gamma parameter, and visualization with Figtree (Rambaut). Significant subclades in the tree are collapsed. In the bottom part of the figure, the consensus sequences of each clade are9given compared with the sequences obtained primate clades. The amino acids that are found in more than 90 % of the sequences are marked by their letter, while the variable regions are represented by an asterisk (”^*^”).

In primates, the IGK exon sequences group into two major clades (Olivieri & Gambon-Deza, 2014). We found that these two IGK clades also exist in rodents, as seen in Figure 3. IGK Clade-I and Clade-II contains the exon sequences of all rodent species included in the study. In Clade II, five subclades exist: the clades IIA, IIB and IIC, which correspond to those of primates (Figure 3 and Table 4, and two rodent-specific subclades, denoted Clade-R1 and Clade-R2, which have no correspondence primate clades. As summarized in Table 4, the great majority of IGK V exon sequences reside in Clades-I, IIC, and the specific rodent clades.

**Table 4:** Distribution of V exons from IGKV and IGLV across clades and species.

|  | IGKV |  |  |  |  |  |  | IGLV |  |  |  |  |
| --- | --- | --- | --- | --- | --- | --- | --- | --- | --- | --- | --- | --- |
| SPECIES | I | IIA | IIB | IIC | mII | IIR1 | IIR2 | I | II | III | IV | V |
| **SQUIRREL-RELATED** |  |  |  |  |  |  |  |  |  |  |  |  |
| *I. tridecemlineatus* | 44 | 2 | 2 | 80 | 0 | 3 | 0 | 11 | 30 | 6 | 4 | 3 |
| **CTENOHYSTRICA** |  |  |  |  |  |  |  |  |  |  |  |  |
| *H. glaber* | 4 | 2 | 2 | 24 | 1 | 1 | 1 | 5 | 4 | 3 | 1 | 0 |
| *C. porcellus* | 25 | 6 | 3 | 81 | 0 | 2 | 0 | 25 | 4 | 8 | 6 | 5 |
| *C. lanigera* | 10 | 11 | 3 | 37 | 1 | 2 | 1 | 15 | 5 | 8 | 1 | 5 |
| *O. degus* | 19 | 1 | 2 | 52 | 1 | 2 | 3 | 12 | 10 | 8 | 4 | 10 |
| **MOUSE-RELATED** |  |  |  |  |  |  |  |  |  |  |  |  |
| *D. ordii* | 26 | 0 | 1 | 59 | 0 | 0 | 0 | 0 | 0 | 0 | 0 | 0 |
| *J. jaculus* | 6 | 3 | 5 | 26 | 0 | 3 | 1 | 0 | 0 | 0 | 0 | 1 |
| *N. galili* | 19 | 2 | 4 | 59 | 0 | 0 | 10 | 6 | 0 | 2 | 2 | 1 |
| *M. ochrogaster* | 4 | 2 | 0 | 37 | 0 | 2 | 2 | 3 | 5 | 0 | 13 | 0 |
| *M. auratus* | 7 | 1 | 6 | 43 | 0 | 6 | 2 | 2 | 4 | 5 | 5 | 0 |
| *C. griseus* | 8 | 2 | 0 | 37 | 0 | 2 | 7 | 0 | 18 | 0 | 5 | 0 |
| *P. maniculatus* | 29 | 3 | 14 | 121 | 0 | 16 | 17 | 3 | 3 | 2 | 4 | 0 |
| *M. musculus* | 16 | 1 | 8 | 49 | 0 | 27 | 10 | 0 | 0 | 2 | 0 | 2 |
| *R. norvegicus* | 56 | 4 | 7 | 81 | 0 | 18 | 10 | 7 | 0 | 2 | 0 | 1 |

Recently, we described the existence of five major evolutionary V gene clades amongst mammals and reptiles in the IGLV locus (Olivieri et al., 2014). Such a shared cladistic structure between species separated by more than 300My of evolution may have originated from functional or mechanistic processes that are still unknown. Despite this longstanding relationship across diverse species, rodents have surprisingly few IGLV genes in comparison. Nonetheless, Table 4 shows that the five major mammal/reptile IGLV clades are conserved in rodents for the families Squirrel-related and Ctenohystrica, but are lost in the Mouse-related families. *D. ordii* does not have lambda chains and *J. jaculus* we has found only one.

### 3.3. V-genes for TRA

In the 14 rodent species studied, we detected 1017 TRA V exons. From our results of sequence annotations (available at http://vgenextractor.org), rodents have the largest number of V genes in the TRA locus as compared to other mammalian orders. We constructed the phylogenetic tree of the AA translated TRAV exon sequences. Similar to the results found in primates (Olivieri & Gambon-Deza, 2014) where we identified 35 evolutionary clades, the TRAV locus in rodents contains multiple clades that group several rodent species. Also, evidence exists that the TRAV locus in rodents underwent recent duplications events. To probe homologous relationships of the TRAV locus between rodents and primates, we constructed a phylogenetic tree consisting of the rodent TRA V exon sequences and the TRA V consensus sequences of the 35 primate clades. The results are shown in Figure 4 and Table 5.

**Figure 4:**
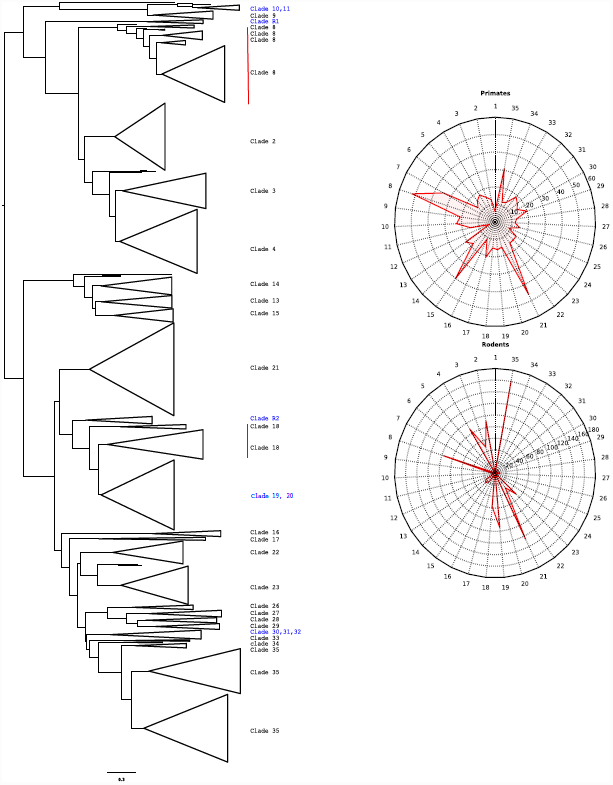
The phylogenetic trees of the AA translated sequences from TRAV exons from 14 rodent species. V exon sequences are obtained from whole genome shotgun (WGS) datasets using the Vgenextractor algorithm. Alignment of the amino acid sequences was performed with clustalO (Sievers & Higgins, 2014), tree construction with FastTree (Price et al., 2010) using the WAG matrix, and visualization with Figtree (Rambaut). In the right part of the full distribution of the V genes of primates TRAV (total 16 species) and rodents (total 14 species) represented in radar plots for each order. 35 spokes represents each one of the clades. These spokes are marked with the number of exons in each clade. The red line connects the marks.

**Figure 5:**
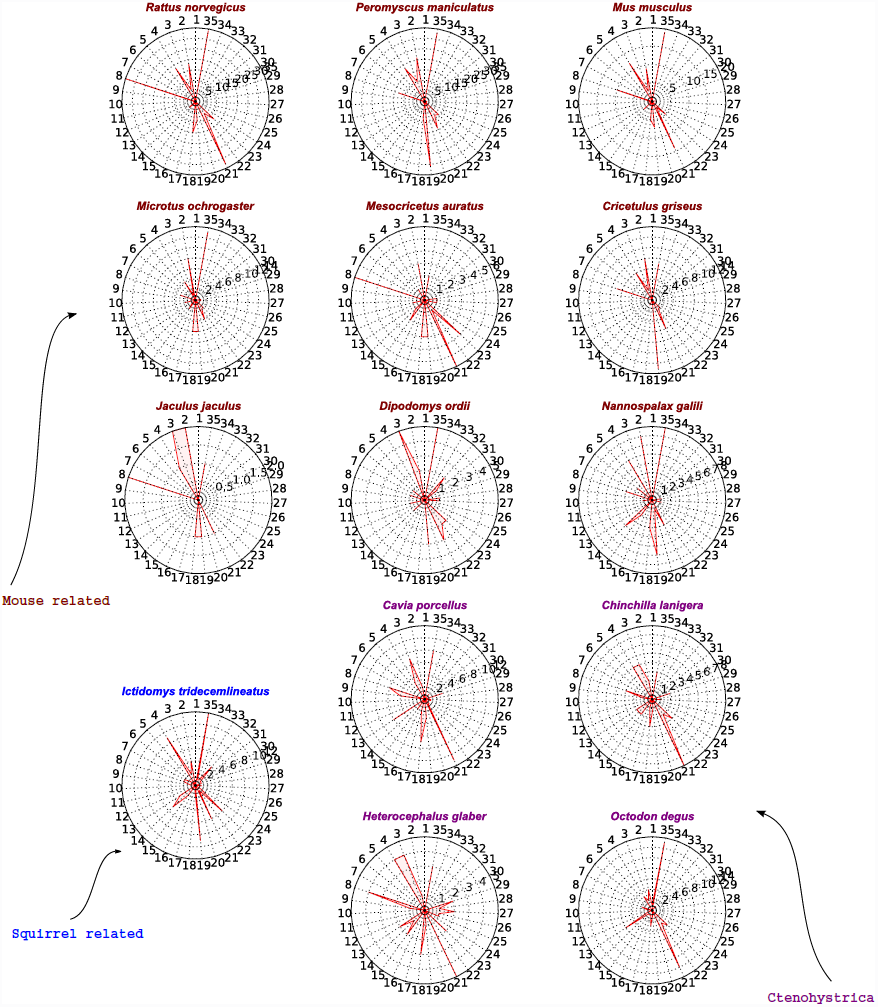
Representation of the number of V exons in the TRA locus of rodents. For each species, circular radar plots, with axes for each of the 35 clades, are used to indicate the number of V exons belonging to each clades.

**Table 5:** Number of TRAV exons present in each clade by specie in the phylogenetic tree defined in Figure 3.

| CLADE | *I. tridecemlineatus* | *H. glaber* | *C. porcellus* | *Ch. lanigera* | *O. degus* | *J. jaculus* | *D. ordii* | *N. galili* | *M. auratus* | *P. maniculatus* | *R. norvegicus* | *M. musculus* | *C. griseus* | *M. ochrogaster* |
| --- | --- | --- | --- | --- | --- | --- | --- | --- | --- | --- | --- | --- | --- | --- |
| 1 | 0 | 0 | 0 | 0 | 1 | 0 | 0 | 0 | 0 | 0 | 0 | 0 | 0 | 0 |
| 2 | 4 | 1 | 2 | 2 | 4 | 2 | 2 | 7 | 3 | 21 | 18 | 9 | 0 | 8 |
| 3 | 2 | 4 | 7 | 4 | 2 | 2 | 5 | 0 | 0 | 10 | 6 | 3 | 2 | 1 |
| 4 | 9 | 4 | 3 | 4 | 3 | 1 | 0 | 5 | 1 | 18 | 18 | 12 | 6 | 4 |
| 5 | 0 | 0 | 0 | 0 | 0 | 0 | 0 | 0 | 0 | 0 | 0 | 0 | 0 | 0 |
| 6 | 1 | 0 | 0 | 0 | 0 | 0 | 1 | 2 | 0 | 0 | 0 | 0 | 0 | 0 |
| 7 | 2 | 0 | 0 | 0 | 0 | 0 | 0 | 0 | 0 | 0 | 0 | 0 | 0 | 0 |
| 8 | 2 | 4 | 6 | 3 | 2 | 2 | 1 | 3 | 6 | 13 | 35 | 10 | 7 | 3 |
| 9 | 1 | 1 | 4 | 1 | 1 | 0 | 0 | 0 | 0 | 1 | 1 | 1 | 0 | 0 |
| 10 | 0 | 0 | 0 | 0 | 0 | 0 | 0 | 1 | 0 | 0 | 0 | 0 | 0 | 0 |
| 11 | 0 | 0 | 0 | 0 | 0 | 0 | 1 | 0 | 1 | 1 | 1 | 1 | 0 | 2 |
| 12 | 0 | 0 | 0 | 0 | 1 | 0 | 0 | 0 | 0 | 0 | 0 | 0 | 0 | 0 |
| 13 | 3 | 2 | 6 | 2 | 6 | 0 | 0 | 0 | 0 | 0 | 0 | 0 | 0 | 0 |
| 14 | 5 | 1 | 0 | 2 | 1 | 0 | 1 | 4 | 1 | 1 | 4 | 3 | 0 | 1 |
| 15 | 2 | 2 | 1 | 2 | 1 | 0 | 0 | 2 | 2 | 1 | 2 | 0 | 0 | 2 |
| 16 | 0 | 1 | 1 | 1 | 1 | 0 | 0 | 0 | 0 | 1 | 1 | 1 | 1 | 0 |
| 17 | 1 | 0 | 0 | 0 | 0 | 0 | 0 | 0 | 0 | 0 | 0 | 0 | 0 | 0 |
| 18 | 1 | 3 | 7 | 3 | 3 | 1 | 0 | 3 | 3 | 8 | 15 | 5 | 0 | 6 |
| 19 | 9 | 1 | 3 | 1 | 1 | 1 | 3 | 6 | 3 | 31 | 9 | 7 | 13 | 6 |
| 20 | 0 | 0 | 0 | 0 | 0 | 0 | 0 | 0 | 0 | 0 | 0 | 0 | 0 | 0 |
| 21 | 6 | 5 | 13 | 9 | 13 | 1 | 3 | 4 | 6 | 14 | 33 | 14 | 6 | 4 |
| 22 | 1 | 0 | 1 | 2 | 3 | 0 | 2 | 0 | 1 | 8 | 7 | 2 | 2 | 2 |
| 23 | 6 | 2 | 1 | 3 | 4 | 0 | 2 | 0 | 2 | 9 | 12 | 5 | 2 | 2 |
| 24 | 1 | 0 | 0 | 0 | 0 | 0 | 0 | 0 | 0 | 0 | 0 | 0 | 0 | 0 |
| 25 | 0 | 1 | 0 | 0 | 0 | 0 | 0 | 0 | 0 | 0 | 0 | 0 | 0 | 0 |
| 26 | 0 | 1 | 1 | 0 | 0 | 0 | 0 | 1 | 1 | 1 | 0 | 0 | 0 | 0 |
| 27 | 1 | 2 | 1 | 1 | 0 | 0 | 1 | 0 | 1 | 1 | 0 | 0 | 0 | 1 |
| 28 | 0 | 1 | 0 | 0 | 0 | 0 | 1 | 1 | 1 | 1 | 0 | 1 | 0 | 1 |
| 29 | 1 | 2 | 3 | 2 | 0 | 0 | 0 | 0 | 0 | 0 | 0 | 0 | 0 | 0 |
| 30 | 1 | 0 | 0 | 0 | 0 | 0 | 0 | 0 | 0 | 0 | 0 | 0 | 0 | 0 |
| 31 | 1 | 0 | 0 | 0 | 0 | 0 | 1 | 0 | 0 | 0 | 0 | 0 | 0 | 0 |
| 32 | 4 | 0 | 1 | 0 | 1 | 0 | 2 | 0 | 0 | 0 | 0 | 0 | 0 | 0 |
| 33 | 1 | 0 | 0 | 0 | 0 | 0 | 0 | 0 | 0 | 0 | 1 | 0 | 0 | 0 |
| 34 | 1 | 0 | 0 | 0 | 0 | 0 | 0 | 0 | 0 | 0 | 0 | 0 | 0 | 0 |
| 35 | 12 | 3 | 8 | 3 | 13 | 1 | 5 | 8 | 2 | 33 | 34 | 19 | 7 | 13 |

From the cladistic relationships of the TRAV locus, we can describe in more detail than previously possible the large increase of V genes in this locus. In particular, the clades 8, 19, 20, 21 and 35 (of Figure 4 underwent an expansion and several subclades were generated during the evolutionary diversification of rodents. Also, approximately one third of the primate TRAV clades are absent in rodents (the clades of Figure 4: 1, 5, 7, 10, 12, 17, 20, 24, 25, 30, 31 and 34). Thus, we can conclude that in rodents, there is an increase in the number TRA V genes that is concentrated within a few clades and that this expansion process has been accompanied with the loss of representatives in other clades. In Figure 4(right), the distribution of primates and rodents TRA V genes is represented. The clade distribution in primates is more homogeneous (i.e., the number of species per clade is approximately constant) than in rodents, supporting the hypothesis of a large inter-order expansion of the rodent TRAV clades.

We also studied the number of genes per clade for each species (5). In the Squirrel-related rodent,*Ictidomys tridecemlineatus*, there is an expansion of V sequences in clades 4, 19, and 35 (i.e., each clade consists of 9, 9 and 12 TRAV genes, respectively). In each of the four Ctenohystrica species clade 21 underwent an expansion. In Mouse-related species, some species exhibit large expansions. For example, the rat has more than 30 members in each of the clades 8, 21 and 35. In other Mouse-related species, the clades that expand are not always the same. In *P. maniculatus*, there is an expansion in clade 35, as in the rat, but additionally, there is also an expansion in clade 19, don’t see in rat. The significance of these V gene expansion and cladistic dependencies is unknown.

### 3.4. TRB V genes

In the rodent species we found 327 V exons in the TRB locus. This locus contains approximately 1/3 the number of V exons found in the TRAV loci. Figure 6 shows the phylogenetic tree of the AA translated V exon sequences. As before, we included the TRBV primate consensus sequences in the alignment to identify orthologous clades. As seen in Table 6 and Figure 6, the number of taxa and sequences per clade is approximately constant, without notable expansions. Clade distribution is more homogeneous than the TRAV locus. The clearest example is seen in *R. norvegicus*, which has the most TRAV gene expansion, but has no more than three sequences in each of the TRBV clades. The TRBV locus in rodents is also characterized by the absence of V genes in clades 10, 15, 17, 18 and 22.

**Figure 6:**
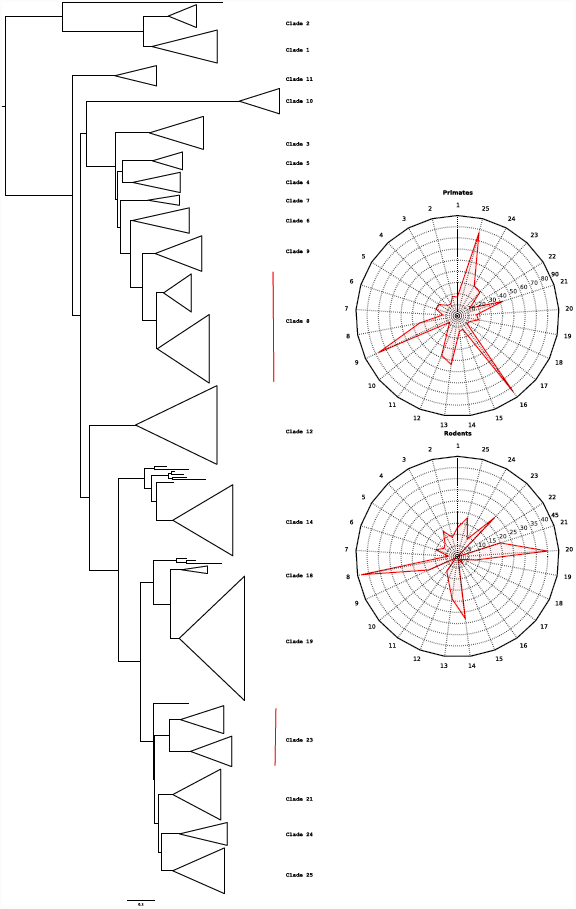
The phylogenetic trees of the AA translated sequences from TRBV exons from 14 rodent species. V exon sequences are obtained from whole genome shotgun (WGS) datasets using the Vgenextractor algorithm. Alignment of the amino acid sequences was performed with clustalO (Sievers & Higgins, 2014), tree construction with FastTree (Price et al., 2010) using the WAG matrix, and visualization with Figtree (Rambaut). (Right) The distribution of TRB V genes of primate (total 16 species) and rodents (total 14 species), represented in circular radar plots for each order, where each spoke represents one of the clades. Spokes serve as axis and they marks the number of exons in each clade. The red line connects these points.

**Table 6:** Number of TRBV exons present in each clade by specie in the phylogenetic tree defined in Figure 3.

| CLADE | *I. tridecemlineatus* | *H. glaber* | *C. porcellus* | *Ch. lanigera* | *O. degus* | *D. ordii* | *J. jaculus* | *N. galili* | *M. ochrogaster* | *M. auratus* | *C. griseus* | *P. maniculatus* | *R. norvegicus* | *M. musculus* |
| --- | --- | --- | --- | --- | --- | --- | --- | --- | --- | --- | --- | --- | --- | --- |
| 1 | 0 | 1 | 2 | 2 | 5 | 0 | 1 | 0 | 1 | 0 | 0 | 0 | 1 | 0 |
| 2 | 1 | 0 | 0 | 0 | 0 | 0 | 0 | 1 | 1 | 1 | 1 | 2 | 1 | 1 |
| 3 | 1 | 1 | 1 | 1 | 2 | 0 | 1 | 1 | 1 | 1 | 1 | 1 | 1 | 0 |
| 4 | 1 | 1 | 1 | 0 | 0 | 1 | 0 | 0 | 1 | 1 | 1 | 1 | 0 | 0 |
| 5 | 1 | 0 | 0 | 0 | 0 | 0 | 0 | 1 | 1 | 0 | 1 | 1 | 1 | 1 |
| 6 | 1 | 2 | 0 | 1 | 0 | 0 | 2 | 0 | 1 | 1 | 1 | 1 | 0 | 0 |
| 7 | 1 | 0 | 1 | 1 | 1 | 0 | 0 | 0 | 0 | 0 | 0 | 0 | 0 | 0 |
| 8 | 3 | 1 | 8 | 1 | 8 | 0 | 1 | 1 | 5 | 2 | 2 | 5 | 3 | 3 |
| 9 | 3 | 1 | 2 | 1 | 1 | 0 | 1 | 1 | 1 | 1 | 1 | 1 | 0 | 0 |
| 10 | 0 | 0 | 0 | 1 | 0 | 0 | 0 | 0 | 0 | 0 | 0 | 0 | 0 | 0 |
| 11 | 1 | 0 | 0 | 0 | 0 | 0 | 0 | 1 | 1 | 1 | 1 | 1 | 1 | 1 |
| 12 | 3 | 0 | 0 | 0 | 0 | 0 | 1 | 1 | 1 | 1 | 1 | 1 | 1 | 1 |
| 13 | 0 | 1 | 3 | 2 | 0 | 0 | 1 | 1 | 1 | 1 | 2 | 4 | 0 | 2 |
| 14 | 2 | 1 | 1 | 1 | 0 | 5 | 1 | 1 | 5 | 2 | 3 | 4 | 2 | 2 |
| 15 | 0 | 0 | 0 | 0 | 1 | 0 | 0 | 0 | 0 | 0 | 0 | 0 | 0 | 0 |
| 16 | 0 | 0 | 2 | 1 | 1 | 0 | 0 | 0 | 0 | 0 | 0 | 0 | 0 | 0 |
| 17 | 0 | 1 | 0 | 0 | 0 | 0 | 1 | 0 | 0 | 0 | 0 | 0 | 0 | 0 |
| 18 | 1 | 0 | 1 | 1 | 0 | 0 | 0 | 0 | 0 | 0 | 0 | 0 | 0 | 0 |
| 19 | 0 | 0 | 0 | 0 | 0 | 0 | 1 | 0 | 1 | 1 | 1 | 1 | 1 | 1 |
| 20 | 1 | 1 | 5 | 4 | 3 | 0 | 4 | 5 | 3 | 2 | 2 | 4 | 3 | 2 |
| 21 | 3 | 1 | 0 | 1 | 0 | 1 | 1 | 1 | 2 | 1 | 2 | 2 | 2 | 2 |
| 22 | 0 | 0 | 0 | 0 | 0 | 0 | 0 | 0 | 0 | 0 | 0 | 0 | 0 | 0 |
| 23 | 4 | 1 | 7 | 2 | 2 | 0 | 1 | 0 | 1 | 1 | 1 | 2 | 1 | 1 |
| 24 | 0 | 0 | 0 | 0 | 0 | 2 | 1 | 1 | 1 | 1 | 0 | 1 | 1 | 1 |
| 25 | 2 | 1 | 4 | 1 | 1 | 2 | 1 | 0 | 2 | 1 | 1 | 2 | 0 | 0 |

**Figure 7:**
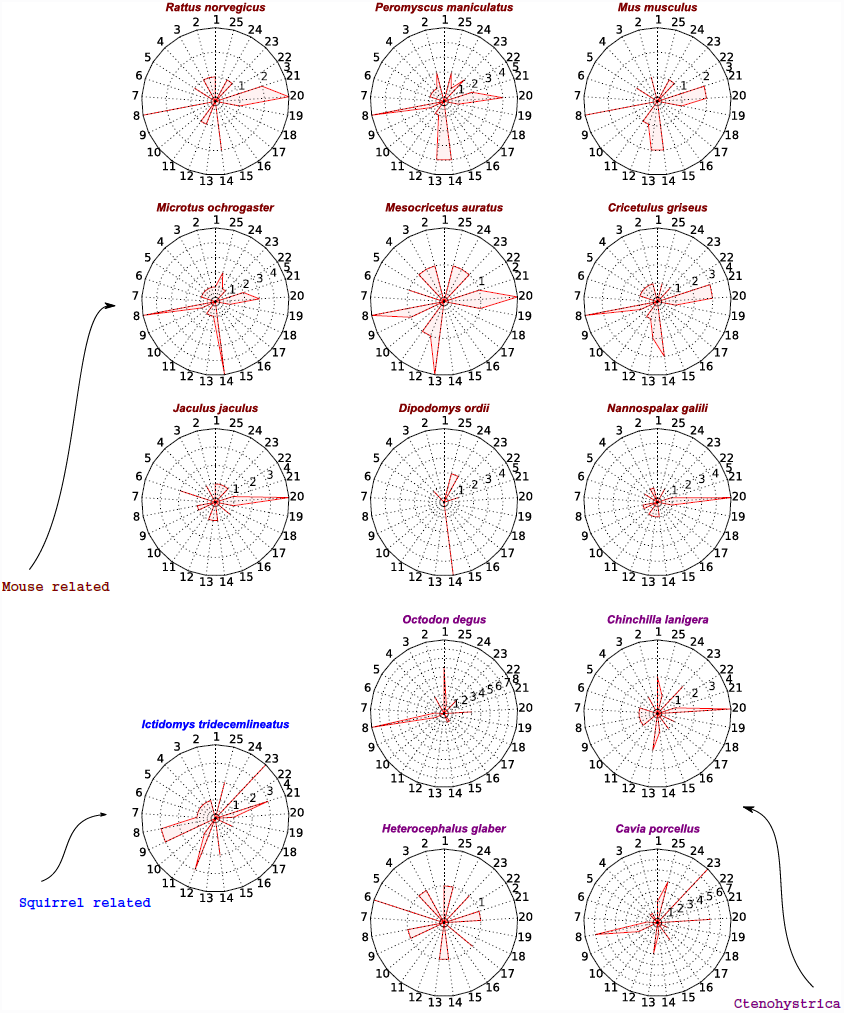
Representation of the number of V exons in the TRB locus of rodents. For each species, circular radar plots, with axes for each of the 25 clades, are used to indicate the number of V exons belonging to each clades.

Figure 7 shows the number of V genes that exists in each clade by species. In this locus, it is rare that a species has more than two genes per clade. There are two exceptional cases: *O. degu* has eight TRB V genes in clade 8, and *C. porcellus* has five and six V genes in the clades 8 and 23, respectively.

### 3.5. Correlations with MHC

Using our random forest based tool to extract MHC genes from WGS datasets (see Methods), we obtained MHC-I and MHC-II (both alpha and beta chains) of rodent species. Table 7 provides a summary of the results and shows the wide variability of MHC-I gene number across species compared to the relative homogeneous distribution for MHC-II. The data for rodent species are plotted in Figure 8. As can be seen a correlation exists between the number of MHC-I genes and the number of genes found in the TRAV locus. In the case of the IG loci, we found a less significant correlation with the number of genes in the IGKV locus, however, no correlation was found between the number of IGHV genes and the number of MHC-I genes. Given these results, we shall carry out more extensive studies of molecular coevolution in the future, following ideas of other work (Lovell & Robertson, 2010; Clark et al., 2012).

**Figure 8:**
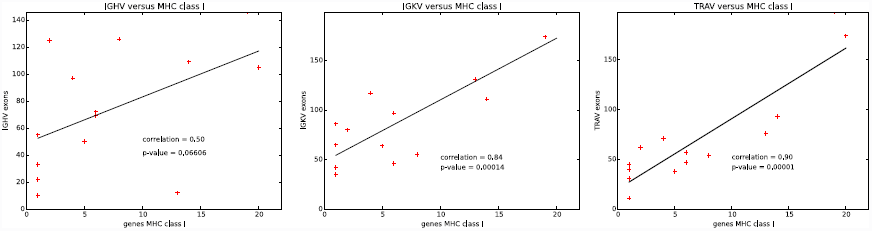
Correlation between the number of MHC class I genes to the number of gene within the loci IGV, IGKV and TRAV. The Pearson test was used to establish the correlation between the two data.

**Table 7:** Number of genes alpha chain of the MHC-I and alpha and beta chains of MHC-II.

| SPECIE | CLASS I | CLASS II A | CLASS II B |
| --- | --- | --- | --- |
| **SKIRREL-RELATED** |  |  |  |
| *I. tridecemlineatus* | 13 | 4 | 5 |
| **CTENOHYSTRICA** |  |  |  |
| *H. glaber* | 1 | 3 | 3 |
| *C. porcellus* | 4 | 2 | 4 |
| *C. lanigera* | 1 | 1 | 4 |
| *O. degus* | 2 | 3 | 6 |
| **MOUSE-RELATED** |  |  |  |
| *D. ordii* | 1 | 0 | 1 |
| *J. jaculus* | 1 | 2 | 3 |
| *N. galili* | 6 | 8 | 4 |
| *M. ochrogaster* | 6 | 3 | 2 |
| *M. auratus* | 5 | 3 | 3 |
| *C. griseus* | 8 | 3 | 13 |
| *P. maniculatus* | 20 | 1 | 3 |
| *M. musculus* | 14 | 2 | 3 |
| *R. norvegicus* | 19 | 3 | 3 |

## 4. Discussion

While the order Rodentia is the mammalian group most proximal in evolution to primates, there is considerable difference between the genomic antigen recognition repertoire structure, particularly amongst the V genes of the IG and TR loci. First, there is a large number of IGHV and IGKV genes, exhibiting a pronounced IGK/IGL ratio. This asymmetry is caused by an expansion of the IGK V gene locus as well as a relative decrease in IGL V genes. The evolutionary mechanisms that have brought about these duplications are elusive, however such differences did occur during the speciation event between rodents and primates, being more pronounced in the mouse-related clade of rodents.

Because the expansion of genes within the IGHV locus occurred in the subclades I-C, II-R, and III-C, the expansion of the IGH locus may not be considered a random process. Moreover, no representative rodent sequences were found in the I-A primate subclade. The data suggests that evolutionary pressures may favor the expansion of particular V genes and facilitate the disappearance of entire subclades. These results suggest that we may expect to encounter such processes in other mammalian orders as well as order specific subclades.

The rodent light chains (i.e., IGK and IGL) are of particular interest. The Mouse-related family of rodents may be the species possessing the largest number of kappa chains (IGKV genes) and fewest lambda chains (IGLV genes). Because of the paucity of IGLV genes, some rodent species lost representation in some of 5 major clans, whose endurance in the mammal and reptile evolution suggests some functional importance. These losses may be the cause or the consequence of the large increase of IGK V genes. Nonetheless, until doubts about the origins and reason for the existence of two light chains are resolved, these results will not be clear. There are other examples of viable species that only possess one of IG light chains. In particular, the absence of IGK chains was described in snakes (Gambó n-Deza et al., 2012) and *M. lucifugus* (Butler et al., 2014). This study of rodents demonstrates that viable species also exist without the IGL light chains.

**Table 8:** Equivalence between the sequences annotated by IMGT and the clades defined in this work

| (a) TRAV |  |  | (b) TRBV |  |  |
| --- | --- | --- | --- | --- | --- |
| CLADE | *H. sapiens* | *M. musculus* | CLADE | *H. sapiens* | *M. musculus* |
| 1 | TRAV40 | — | 1 | TRBV20 | TRBV20 |
| 2 | TRAV18 | TRAV12 | 2 | TRBV29 | TRBV30 |
| 3 | TRAV9-1 | TRAV6 | 3 | TRBV19 | TRBV19 |
| 4 | TRAV9-2 | TRAV6 | 4 | TRBV27 | — |
| 5 | TRAV3 | — | 5 | TRBV28 | TRBV29 |
| 6 | TRAV16 | — | 6 | TRBV25 | — |
| 7 | TRAV8 | — | 7 | TRBV24 | — |
| 8 | TRAV8 | TRAV17,TRAV9 | 8 | TRBV10 | TRBV13 |
| 9 | TRAV4 | TRAV2 | 9 | TRBV6 | TRBV8 |
| 10 | TRAV26-1 | — | 10 | TRBV30 | TRBV31 |
| 11 | TRAV26-2 | TRAV21 | 11 | TRBV15 | TRBV17 |
| 12 | — | TRAV15 | 12 | TRBV4 | TRBV5 |
| 13 | TRAV14 | — | 13 | TRBV4 | TRBV5 |
| 14 | TRAV19 | TRAV16 | 14 | TRBV13 | TRBV12 |
| 15 | TRAV38 | — | 15 | TRAV9 | — |
| 16 | TRAV1 | TRAV1 | 16 | TRBV15 | — |
| 17 | TRAV21,11 | — | 17 | TRBV18 | — |
| 18 | TRAV5 | TRAV3 | 18 | TRBV16 | — |
| 19 | TRAV13-1 | TRAV5,10 | 19 | TRBV21 | TRBV21 |
| 20 | TRAV13-2 | TRAV20 | 20 | TRBV23 | TRBV23,24,26 |
| 21 | TRAV12,29 | TRAV7,14,19 | 21 | TRBV12 | TRBV15,16 |
| 22 | TRAV10 | TRAV11 | 22 | TRBV14 | — |
| 23 | TRAV17 | TRAV8 | 23 | TRBV2 | TRBV3 |
| 24 | TRAV6 | — | 24 | TRBV11 | TRBV14 |
| 25 | TRAV30 | — | 25 | TRBV7 | TRBV9 |
| 26 | TRAV34 | — |  |  |  |
| 27 | TRAV35 | — |  |  |  |
| 28 | TRAV27 | TRAV18 |  |  |  |
| 29 | TRAV25 | — |  |  |  |
| 30 | TRAV24 | — |  |  |  |
| 31 | TRAV36 | — |  |  |  |
| 32 | TRAV20 | — |  |  |  |
| 33 | TRAV39 | — |  |  |  |
| 34 | TRAV41 | — |  |  |  |
| 35 | TRAV22 | TRAV4,13 |  |  |  |

In a previous article, we described orthology between the TR V genes of loci in primates. We studied 16 primate species and identified 35 TRA V genes and 25 TRB V genes that are positively selected. Table 8 shows the equivalence between the annotated IMGT clade nomenclature and the TRAV and TRBV clades defined in this work. Given the evolutionary proximity between the primate and rodent orders, we searched for clade orthology by aligning all the TRAV rodent sequences with consensus sequences derived from each of the major primate TRAV clades. We showed that V gene orthologs do exist in rodents. Moreover, the V gene clades found in rodents correspond to clades found in primates, indicative of their common origin. Such results may suggest that positive selective processes exist in the TR V loci that constrain the number of V genes.

Few major changes are observed in the TRBV locus between primates and rodents. Across the 25 TRBV clades, sequences from species from both orders are evenly distributed, possessing at least one sequence per clade. Such a similar cladistic structuring is not observed in the TRAV locus. In primates, there is a homogeneous distribution and structure similar to that observed in the TRBV clades, having multiple clades but with a few representative sequences per specie within each clade. In rodents, however, there are prominent expansions in specific clades. In *R. norvegicus*, for example, there are more than 25 TRA V genes in each of the clades 8, 21, and 35. A similar situations occurs in other rodent species. This data is interesting because it shows a clear indication that selective evolutionary pressures condition the expansion of the TRAV gene locus without the need for expansion in the TRBV locus. This is an unexpected discovery, because the TRAV and TRBV form heterodimer products and both are involved in antigen-MHC recognition. The fact that there is expansion in the TRAV locus without an expansion in the TRBV locus indicates that that there must be a functional division between the two chains.

Given that both molecules derived from TRA and TRB genes recognize antigens in MHC, the correlation between the number of TRA V genes and the number of MHC class I genes is of great interest. Within V exons, CDR1 and CDR2 are the contact regions during interaction with the antigen-MHC complex. CDR3 however, is generated by somatic processes and is more strongly associated with the recognition of antigen than with the MHC molecule. Thus, coevolution is more likely between the V exons (containing CDR1 and CDR2) and MHC molecule (Housset et al., 1997; Madden, 1995; Brown et al., 1993). Our results in this study suggest that in the evolutionary process in rodent species, such as the rat, mouse and deer mouse, there have been specific concomitant duplication in the TRA V genes and MHC class I genes. While the cause of these duplications requires additional information, one explanation could be due to involute pressure of intracellular infectious agents, considering the fact that the MHC class I presents cytosolic antigens. Also, this coevolution suggests that certain V genes of the TRAV locus (the ones that expanded in these species) are directed towards antigen recognition in MHC class I molecules.

The results presented in this work propose an evolutionary classification of TRA and TRB V-genes in mammals. Such classification may be of interest because it can be used to compare genes between species to uncover their underlying function.

